# Predicting 3D RNA structure from solely the nucleotide sequence using Euclidean distance neural networks

**DOI:** 10.1101/2022.05.16.492153

**Authors:** Congzhou M. Sha, Jian Wang, Nikolay V. Dokholyan

## Abstract

Fast and accurate 3D RNA structure prediction remains a major challenge in structural biology, mostly due to the size and flexibility of RNA molecules, as well as lack of diverse experimentally determined structures of RNA molecules. Unlike DNA structure, RNA structure is far less constrained by base pair hydrogen bonding, resulting in an explosion of potential stable states. Here, we propose a convolutional neural network which predicts all pairwise distances between residues in an RNA, using a recently described smooth parametrization of Euclidean distance matrices. We achieve high accuracy predictions on RNAs up to 100 nucleotides in length in fractions of a second, a factor of 10^7^ faster than existing molecular dynamics-based methods. We also convert our coarse-grained machine learning output into an all-atom model using discrete molecular dynamics with constraints. Our proposed computational pipeline accurately predicts all-atom RNA models solely from the nucleotide sequence.

## INTRODUCTION

Ribonucleic acids (RNA) occupy a central role in biology. In most organisms, they carry information (mitochondrial RNA) and construct proteins (transfer RNA and ribosomal RNA). RNAs can also perform specialized functions such as chemical sensing (riboswitches) and enzymatic catalysis (ribozymes) (1). While 2-3% of the human genome is transcribed in mRNA for protein production, recent estimates suggest that as much as 70-80% of the human genome is transcribed into RNAs with unknown function, the so-called “dark matter” non-coding RNA (2–4). The diversity of structure and function RNA possesses have led to the RNA world hypothesis, in which life on Earth is proposed to have started with RNA performing all the functions that proteins have since taken over (5). Recently, researchers have started engineering RNAs to perform specific functions, from basic science applications, such as genome engineering (6) and protein labeling (7), to clinical applications, such as vaccines (8, 9), therapeutics (10), and diagnostics (11). The common thread in these applications is the need for an accurate 3D RNA structure.

Here, we use state-of-the-art neural network architectures and symmetries to create a fast and accurate neural network which predicts the 3D structure of an RNA solely from its nucleotide sequence. We call our approach the Euclidean Parametrization of RNA (epRNA).

While one might expect that highly successful existing techniques for protein structure prediction may be applicable to RNA structure prediction, there are key differences between proteins and RNAs which present significant obstacles to this transfer of knowledge. First, each residue in a protein has a well-defined hydrophobic/hydrophilic character and size (12). These physical aspects significantly constrain the energy landscape of a protein, so that hydrophilic residues are typically located near the surface of the protein whereas hydrophobic residues are buried in the protein interior. In contrast, the four canonical nucleotides in RNA (adenine, cytosine, guanine, uracil) have far less diversity in hydrophobic character or size, providing less driving force per residue to fold into a specific conformation. Second, each protein residue can be characterized by three backbone dihedral angles and one side chain dihedral angle (13), with the exceptions of glycine and proline. As a result, the three backbone dihedrals per residue largely determine the protein’s 3D structure, and constraints exist on the most likely combinations of dihedral angles (14). In contrast, the phosphate-ribose backbone of RNA has six backbone dihedrals per nucleotide (15), significantly increasing the number of conformations that an RNA of a given length can adopt in comparison to a protein of the same length. Third, far fewer RNA structures are known (~10^3^) in comparison to proteins (~10^5^) (16). This paucity of known structures may be explained as a combination of the factors discussed above: a given RNA may not adopt a single stable conformation due to the lack of large energetic biases and due to the large conformational sample space.

Many existing methods for 3D RNA prediction involve molecular dynamics simulations, sequence alignments, and additional human expert knowledge (17–27). The force field parameters used in RNA molecular dynamics are typically fitted using experimental data (28, 29). A recently published effort to incorporate machine learning with geometric symmetries into the prediction pipeline fails to generate any 3D structures (30).

Here, we overcome the computational cost of molecular dynamics-based RNA folding methods and address the inability of existing neural network models to produce 3D coordinates for RNA. We use a parametrization of Euclidean distance matrices introduced by Hoffmann and Noé (31) to enable the neural network to directly output all inter-residue distances, which we subsequently convert into an all-atom RNA structure using discrete molecular dynamics with constraints. Our computational pipeline is fast and accurate, and we demonstrate its performance on RNAs up to 67 nts in length.

## METHODS AND MATERIALS

### Data collection

We search the Protein Data Bank (PDB) for RNA-containing structures for which the refinement resolution was less than 5 angstroms (median 2.4 angstroms, interquartile range: 1.05 angstroms, see Supplemental Figure 1), the Polymer Entity Type was equal to RNA and none of the other categories (DNA, Protein, NA-hybrid, Other), and the Number of Distinct RNA Entities was equal to 1. As of March 26, 2022, the results totaled 788 structures. We download the .pdb files for these structures and extracted atomic coordinates. We split the structures randomly into training and test sets in a 60:40 ratio.

### Data preprocessing for neural network

We then group the atoms by nucleotide and found the unweighted centroid of each group of atoms, calculating the resulting pairwise distance matrix. We also store the nucleotide sequence in the order present in the .pdb file as a one-hot encoded vector (A, C, G, U, DA, DC, DG, I, N). DA, DC, and DG stand for the 2’-deoxy versions of the RNA bases; I stands for inosine; N stands for ambiguous nucleotide.

### Convolutional neural network

We implement neural networks in TensorFlow 2.8.0 (32) (Figure 1). The specific network we used had 1,257,345 trainable parameters. To convert the one-hot encoding *S* as a square matrix *M*, we perform a Kronecker product, a tensor product in the one-hot encoding basis, where ~ indicates isomorphism:

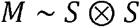

In terms of dimensions, if the sequence has length *s* and the one-hot encoding has *c* channels, the output tensor is of shape (*n, n, c, c*). We then flatten the last two dimensions of the array, organizing the entries into a single *c × c* dimension, and *M* may be interpreted as an *n × n* matrix for which the ordered pair of nucleotides is represented by 1 in a specific channel and zeros elsewhere in the last dimension, which has length *c*^2^. We did not symmetrize or antisymmetrize the tensor product since there is an intrinsic orientation in RNA from 3’-hydroxy to 5’-hydroxy.

**Figure 1:**
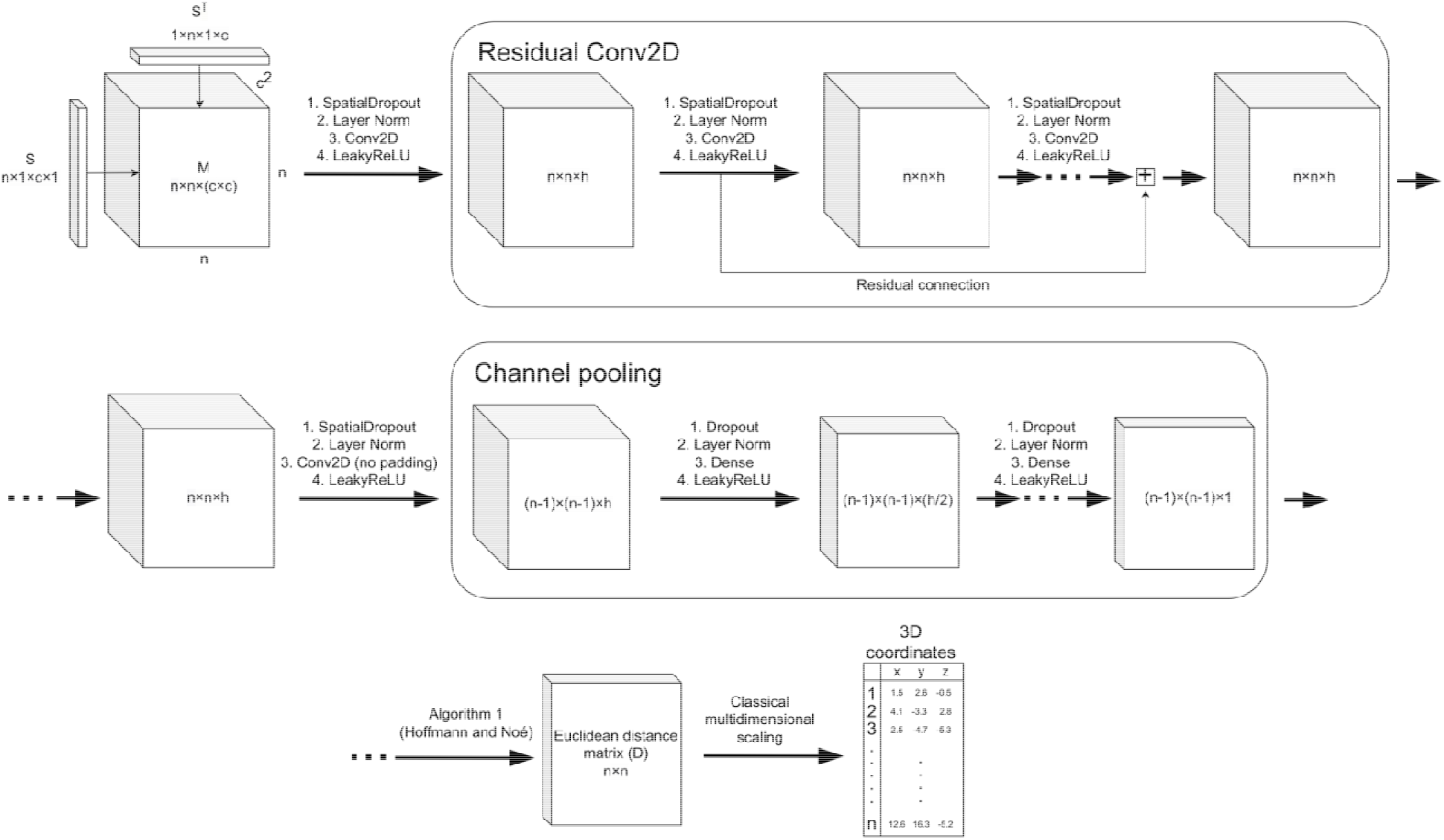
Pipeline for generating Euclidean distance matrices and 3D coordinates purely from an RNA sequence. In the first row, is the one-hot encoding with levels of an RNA sequence of length. We represent all the pairwise interactions of nucleotides by forming a tensor product of with itself (), called. We coerce the data into an array shape amenable for two-dimensional convolutions. In the Residual Conv2D block of the network, we apply standard 2D convolutions (with padding) common in image processing; we use spatial dropout for regularization, layer normalization and residual connections to prevent vanishing and exploding gradients, and LeakyReLU activations. In the second row, we perform one final convolution with no padding, and use fully connected layers to reduce the intermediate representation into the form required for Algorithm 1. The Channel Pooling block is also known as a 1×1 convolution. In the final row, we apply Algorithm 1 to obtain the Euclidean distance matrix for the RNA sequence and convert the distances into 3D coordinates using the distance geometry algorithm known as classical multidimensional scaling (MDS). Algorithm 1 and MDS are smoothly differentiable operations, therefore we can propagate derivatives backward either through RMSD in the 3D coordinates or through a direct comparison of the generated Euclidean distance matrix to the ground truth. Since all layers in this network do not explicitly require a certain value of *n*, we are able to use the same weights for RNAs of arbitrary length, both in training and inference.

We then process *M* through convolutional layers with padding of the same values along the borders (32, 33), keeping the intermediate matrices the same size. We also apply the LeakyReLU activation function with leak rate 0.3 (34). There was no padding in the last convolutional layer, so that the result was an (*n* − 1) × (*n* − 1) matrix, as required by the EDM parametrization. We apply residual connections (35) and 2D spatial dropout (36) throughout the convolutional part of the network. Next, we process the result of the convolutional layers through a series of dense layers, from *c* channels down to 1 channel. We apply dropout in the dense layers (37). We also use layer normalization throughout the model, except for the last dense layer (38). We apply Algorithm 1 from Hoffmann and Noé to the output (31). We use the Adam optimizer with a learning rate of 0.001 (39). Training was carried out on a single NVIDIA Tesla T4 GPU (16 GB VRAM) managed by a single core of a Xeon E5-2680 processor, and training to 100,000 epochs of the neural network implemented in our code completed within 48 hours. Reasonable RNA predictions were available at 10,000 epochs.

### Generating valid Euclidean distance matrices and the corresponding 3D coordinates

Here, we reproduce Algorithm 1 from Hoffmann and Noé (31). Given an arbitrary (*n* − 1) × (*n* − 1) matrix such as that produced by our neural network *A = model(S)*, we symmetrize the matrix

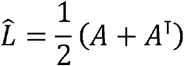

We make the matrix positive using an elementwise softplus activation function

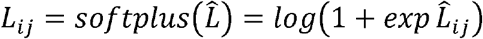

As noted in the original paper, there are alternatives activation functions to softplus. Next, we organize *L* into a Gram matrix *M*

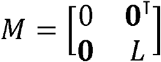

where **0** is column of zeroes with length *n* − 1. Finally, we create a matrix *D* (which is not necessarily a Euclidean distance matrix in 3D),

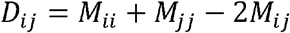

To ensure that the distance matrix is appropriate for three-dimensional Euclidean space, we first calculate

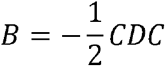

where 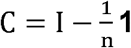 and **1** is an *n × n* matrix of all ones. *B* is known as the strain in multidimensional scaling (40), and serves two purposes here.

First, *D* is a Euclidean distance matrix if and only if *B* is positive semidefinite (41), i.e. the eigenvalues are all nonnegative. To ensure that *B* corresponding to our model output is positive semidefinite, we add a loss on the negative eigenvalues *λ_k_* of *B* (31)

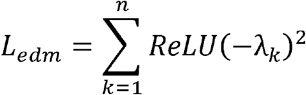

where the rectified linear unit function is also known as the ramp function, and is defined as *ReLU*(*x*) = *max*(0, *x*).

Second, we can generate 3D coordinates from *B*. We compute the *d* = 3 largest eigenvalues and eigenvectors of *B*. The coordinates are

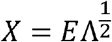

where *E* is the matrix of the largest *d* eigenvectors as columns, Λ is the *d* × *d* matrix of the corresponding eigenvalues along the diagonal and zeros elsewhere, and the rows of *X* are the output coordinates.

If the distance matrix corresponds to the Euclidean metric between points located in 3D space, then all other eigenvalues will be zero (31). Therefore, to ensure that the dimension is 3, we add a loss on all other eigenvalues

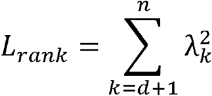

where *λ_k_* ≥ *λ*_*k*+1_.

We add *L_edm_* and *L_rank_* to our training loss with an overall factor of 1.

### Loss function and regularization

We train the network on the elementwise squared difference between the neural network output *model*(*S*) and the true distance matrix *D_true_*

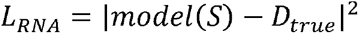

We enforce the dimension of the EDM parametrization to be equal to *d* = 3 through additional losses described above. We regularize all parameters *p_i_* with squared norm with a relative weight of 0.01:

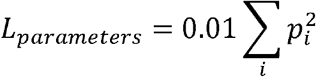

We used a dropout rate of 0.2. Our overall cost function is

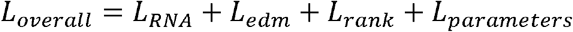

### Alignment of 3D point clouds

We used the well-known Kabsch algorithm (42) to align structures, without the correction for chirality of the best fit rotation.

Given sequences of vectors which are centered so that the centroid of each point cloud is at the origin, we arrange the vectors into matrices *X* and *Y*, in which the columns of the matrices are the components of each vector and the rows in the matrices correspond to pairs of points which are to be matched, one computes the covariance matrix

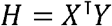

and then the singular value decomposition

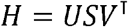

The best fit rotation (and possible reflection) from the vectors in *X* to *Y* is given by

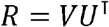

We did not include the correction for chirality, which involves negating one of the coordinates in *X*.

After alignment, we calculated the root mean square deviation (RMSD) between the aligned sequences of vectors {*x*_1_, *x*_2_, ⋯} and {*y*_1_, *y*_2_, ⋯} to compute 3D structure similarity:

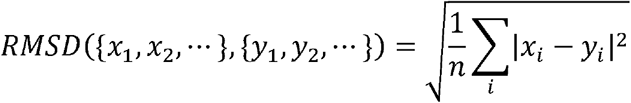

### Clustering of primary sequences

To compute an approximate similarity between two RNAs, we calculated the Levenshtein edit distance *L*(*i*, *j*) between two sequences (43) and divided by the minimum of the two RNA lengths:

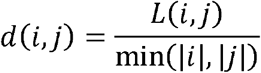

We performed hierarchical clustering of our RNA dataset with scikit-learn 1.0.2 (44), using a dissimilarity cutoff of 0.4.

### Molecular dynamics simulations

We employed discrete molecular dynamics (DMD) (45–47) with an RNA force field and constraints from epRNA model outputs to further refine structures as well as create an all-atom model of the RNA (48, 49). For DMD, we used no additional information other than the primary sequence and the distance constraints from a trained epRNA model. For all distances predicted by epRNA to be under 8 angstroms, we placed a constraint of that distance +/− 2 angstroms on the C3’ atom in the backbone. We clustered the lowest 1% energy structures, with a clustering cutoff of 15 angstroms, and calculated RMSD to the crystal structure.

For coarse-grained RNA folding, we used the Nucleic Acid Simulation Tool (NAST) (25), performing simulations for several thousand iterations at 300 K. The starting conformation was the epRNA predicted coordinates, and we took the final conformation from NAST as the predicted output.

### RNA structural prediction significance

We used a tool by Hajdin et al. (50) to estimate the statistical significance of an RNA structural prediction as a function of nucleotide length and RMSD to the true structure. We report p-values for structures determined without supplying the true secondary structure as input.

## RESULTS

### Architecture and training

We create a convolutional neural network (35) with Euclidean distance parametrization (Figure 1), incorporating state-of-the-art components for efficient training, inference, and data usage. The network contains approximately 10 layers and 1.2 million trainable parameters for all our experiments, and did not perform hyperparameter optimization (e.g. number of layers, filter size, learning rate). We train our networks to minimize the loss described in **Methods**. The primary objective was to minimize the squared difference between the model’s output distance matrix and the ground truth distance matrix.

### Accuracy

In a uniformly random split of the RNAs into training and validation sets, we achieve statistically significant root mean square deviations (RMSDs) on our validation set (Figure 2). To evaluate the bias introduced by correlations among our data, we perform multiple cross-validation studies. First, we hierarchically cluster the RNAs using a measure of distance between nucleotide sequences as described in **Methods** (Figure 3A). We left out the largest cluster from a training set, train a model on the remaining structures, and evaluate the model on the largest cluster (Figure 3B). Second, we train only on RNAs less than 50 nts in length, evaluating on the remaining RNAs (Figure 3C). Third, we train only on RNAs between 50 and 100 nts in length, and evaluating on the remaining RNAs (Figure 3D). In all our cross-validation studies, the neural networks successfully learned the training data and generalized to the validation data.

**Figure 2:**
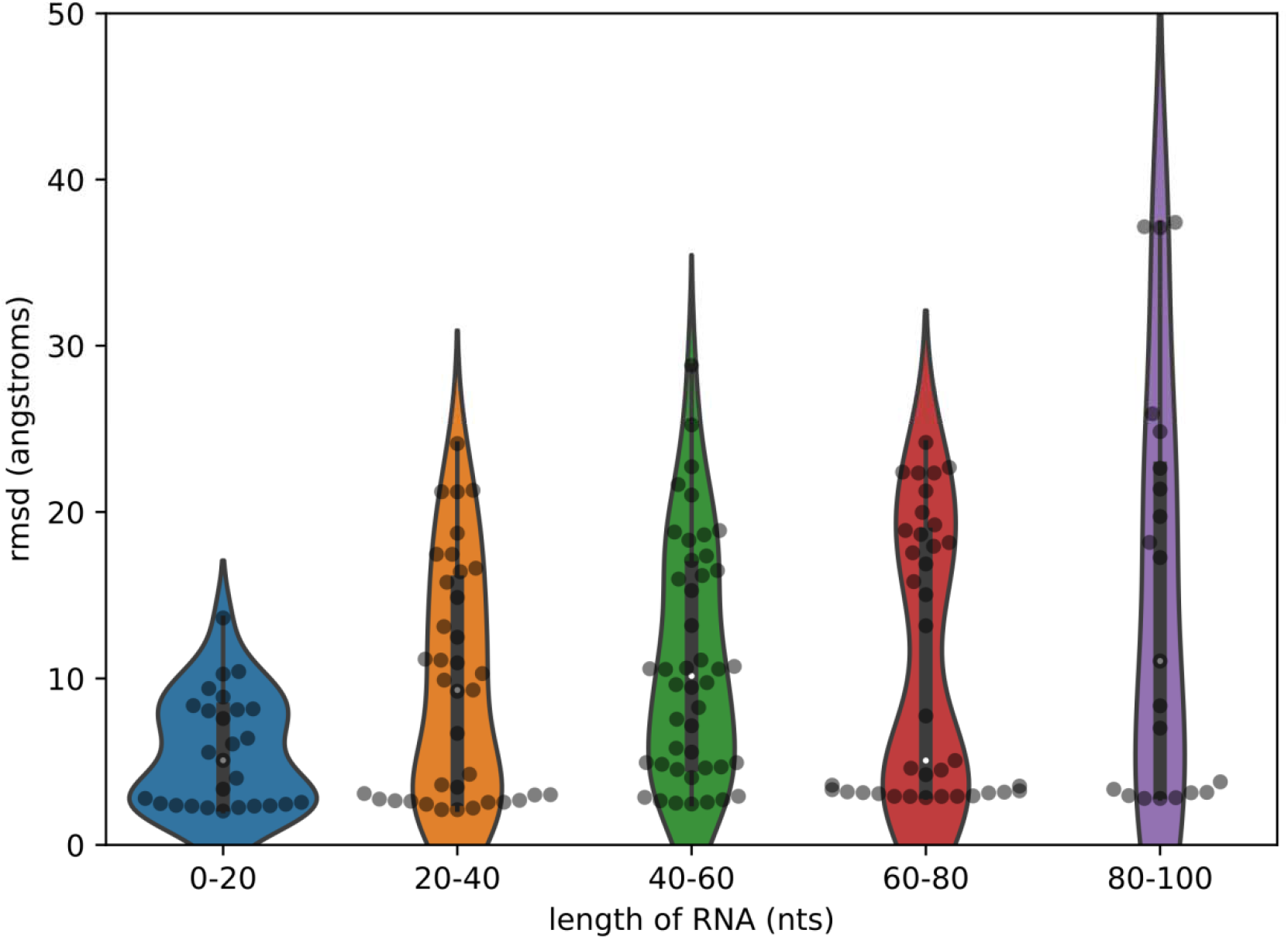
The result of training an epRNA model on a 70:30 random split of the data into a training set (n=352) and validation set (n=226). The accuracy (RMSD) of epRNA models on the validation set is shown, which is stratified by number of nucleotides in the RNA. We omit RNAs in our dataset longer than 100 nts and shorter than 6 nts. We achieve reasonable performance on nucleotides of varying lengths.

**Figure 3:**
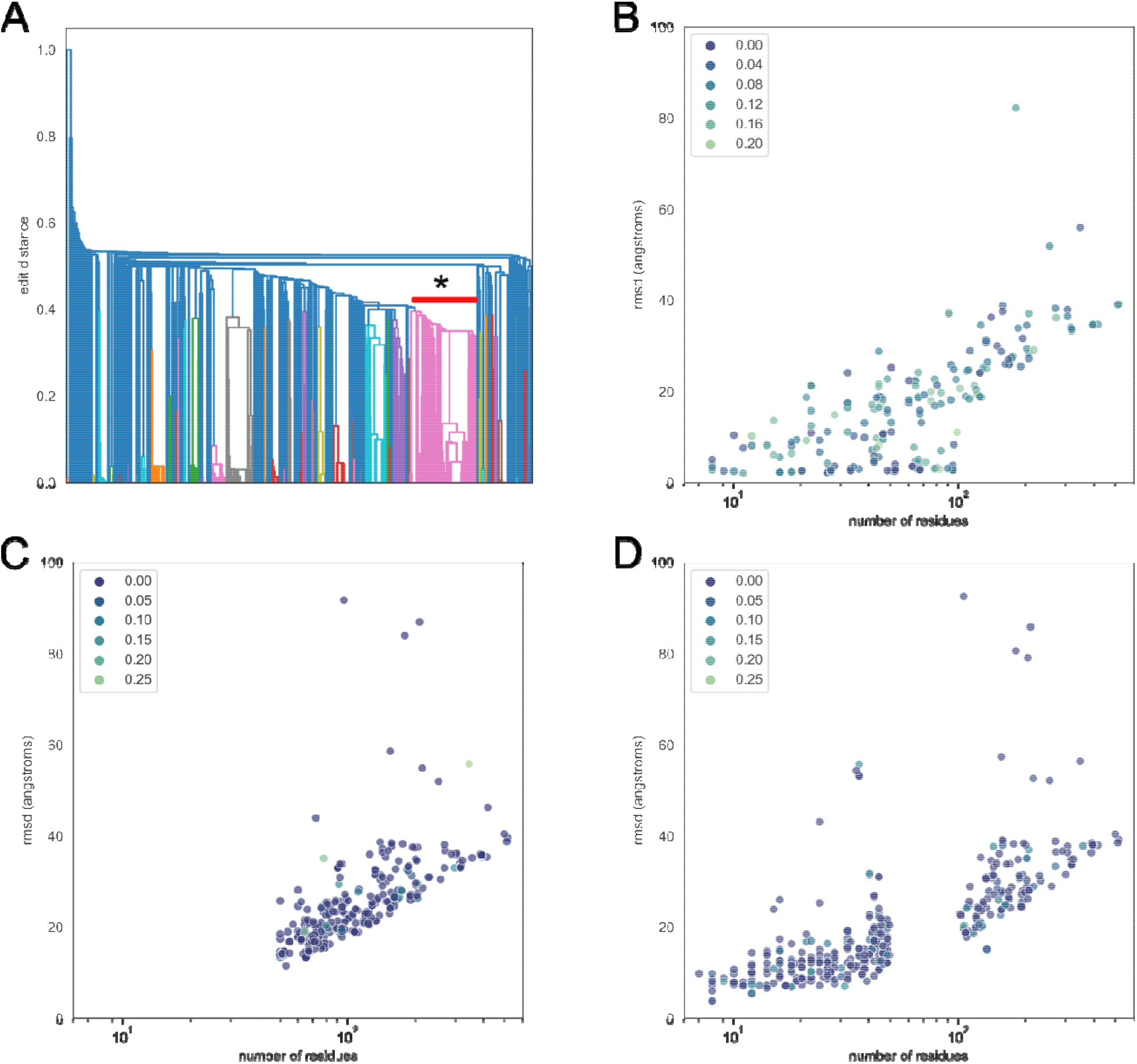
Multiple cross-validation studies for the generalizability of the epRNA approach. **A**: Hierarchical clustering of RNA structures by Levenshtein edit distance, with larger edit distances indicating higher dissimilarity. We took the largest cluster (*) when imposing a cutoff of 0.4 for dissimilarity for leave-one-out validation. **B**: Leave-one-out validation, in which we trained on all single-chain RNAs under 100 nts in length which were not in the largest cluster (*) from **A**, and subsequently evaluated RMSDs on (*). **C**: Short nucleotide validation, in which we trained on RNAs under 50 nts in length, and evaluated RMSD on RNAs greater than 50 nts in length. **D**: Long nucleotide validation, in which we trained on RNAs with length, and evaluated on nucleotides with length outside that range. Each point in **B, C, and D** represents a structure in the respective validation set. The largest structures in the validation sets were approximately 500 nts long. We evaluated the number of structures in the respective training sets which were within an edit distance of 0.4 of each point, and expressed this number as a percentage of the training set. For example, a point with a color corresponding to a value of 0.05 was similar to approximately 5% of the training set, with an edit distance cutoff of 0.4. We omitted RNAs longer than 100 nts for training efficiency purposes, and greater than 600 nts for technical reasons, as the intermediate matrices grow in memory as *n*^2^. The x-axes (number of residues) are plotted on a log scale, and both axes are shared among **B, C, and D** for easy interpretation.

### Speed

Inference on an NVIDIA Tesla T4 averaged approximately 40 sequences per second, when performed on all the structures available. Compared to typical discrete molecular dynamics simulations (DMD) (conservatively 1 day (45–49)), we achieved nearly 10^7^ faster determination of structure. Furthermore, a single molecular structure is returned, and no clustering of molecular trajectories or ensembles is needed.

### Further molecular force field refinement of RNAs

Using the results of the leave-one-out analysis (Figure 3B), we attempted to refine our structures and convert to full-atom models using constraints, both in a coarse-grained force field model of RNA (25) and in a DMD (45–49) setting. We found that further molecular simulation resulted in increased RMSD for all seven structures we examined (Supplemental Figures 2 and 3, Supplemental Table 1), except for one 67 nucleotide structure (Figure 4). In that structure (PDB ID 4FEO), DMD was able to improve the RMSD from 10 angstroms to 7.1 angstroms. Most predicted structures achieved statistical significance (see **Methods**) at the level of p<0.05, with many having p<0.001.

**Figure 4:**
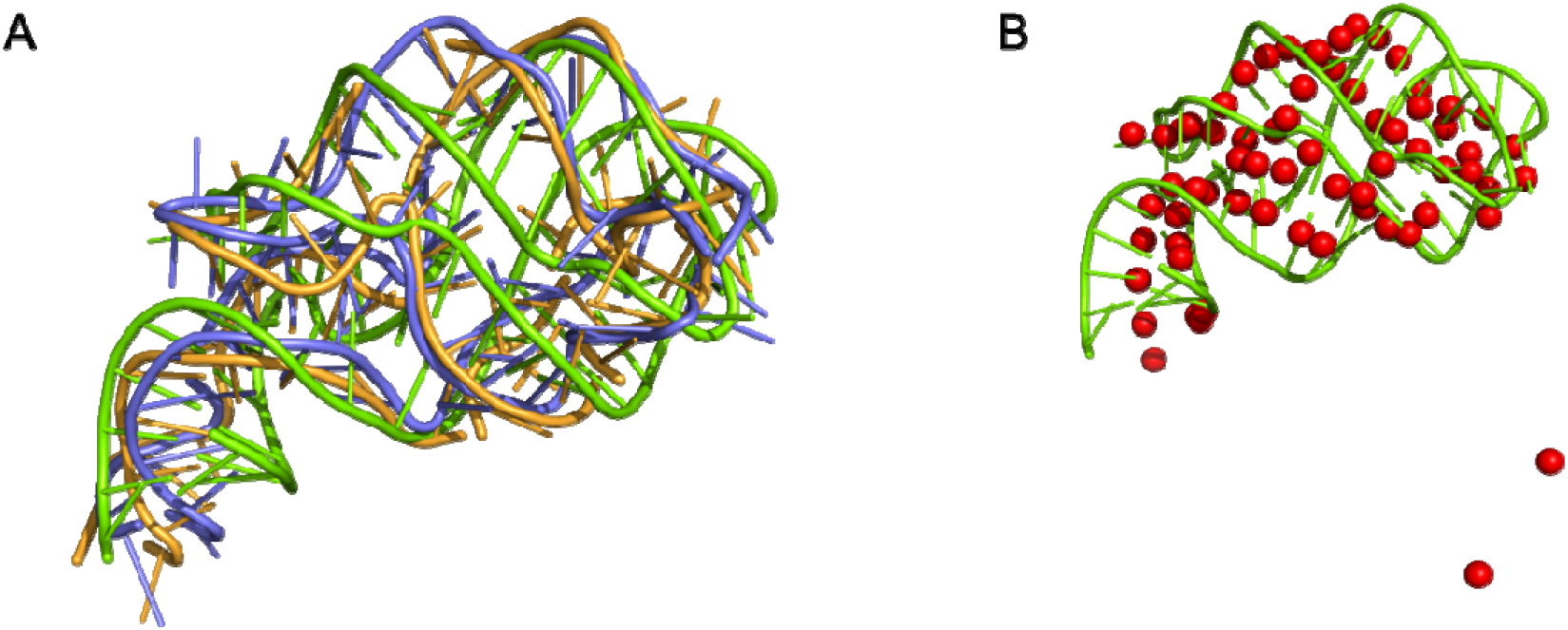
PDB ID 4FEO, a single-chain RNA with 67 nucleotides, along with full-atom models obtained through constrained discrete molecular dynamics. **A**: Green represents the true structure, orange (7.2 angstrom RMSD, p<0.001) and purple (7.6 angstrom RMSD, p<0.001) represent structures obtained from clustering of molecular trajectories obtained from discrete molecular dynamics using the epRNA’s output as distance constraints. P-values are the probabilities of predicting a 67 nucleotide RNA structure with the given RMSD by chance. **B**: Green is the true structure, while red are the epRNA predicted backbone coordinates (10 angstrom RMSD). For this structure, discrete molecular dynamics was able to fix the two outliers in the bottom right of **B** without disrupting the rest of the structure. When we performed a coarse-grained simulation (NAST) for this purpose, the RNA was severely misfolded, resulting in an RMSD of greater than 18 angstroms (Supplemental Figure 2).

### Size dependence

We found a sublinear increase in RMSD as a function of nucleotide length (Figures 3B, 3C, 3D), which may even be logarithmic. Unfortunately, we believe there are too few structures in the Protein Data Bank for strong conclusions to be drawn: there may be entire families of RNAs which are not represented in known structures. There are also likely RNAs which do not have well-defined structures and are, instead, intrinsically disordered, just as are some proteins (51).

## DISCUSSION

Our epRNA approach is robust across various sizes of RNAs (Supplemental Table 1) and demonstrates generalizes to other structures in the cross-validation studies that we performed (Figure 3). Additionally, our method is rapid and highly data efficient, requiring solely the sequence of an RNA of arbitrary length. Performing molecular dynamics simulations did not significantly improve RMSDs for most structures, however the simulations enabled creation of all-atom models from our coarse-grained output.

### Choice of architecture

We hypothesize that just as the base pairing matrix viewed as an image has local structure, so too does the distance matrix. For those residues which are base paired or otherwise physically close, the surrounding residues are also more likely to interact with each other. Due to this reasoning, we treated the input interaction matrix as an image and used 2D convolutional layers to compute intermediate features.

Because 3D RNA structure is invariant under rotations and translations of coordinate frame, we hypothesized that by producing an output invariant under the special 3D Euclidean group, SE(3) (52, 53), we could create a strong inductive bias toward predicting RNA structures. Our application of symmetry was especially important since so few known RNA structures exist and producing a symmetry-invariant output would significantly reduce the number of parameters needed in such a model, reducing the likelihood of overfitting and the need for artificial data augmentation.

We use the E(3)-invariant EDM parametrization rather than alternative methods which achieve SE(3)-equivariance (54), a distinction explained in the following section.

### The role of symmetries in deep learning

Leveraging symmetries to simplify a mathematical problem typically produces significant insight into the problem, as well as increasing solution efficiency (55). In the context of neural network engineering, symmetries reduce the number of parameters necessary to model a function (56), improving both the generalizability of the model and the computational speed of the model.

The most basic example of symmetry in neural network engineering is the convolutional layer (57), in which an input is hypothesized to experience a symmetry under translation. The symmetry in convolutions is **equivariant**, in which a transformation on the input causes the same transformation on the output. The symmetry for a 2D convolutional is termed an T(2) equivariance, in which the symmetry is described by translations in the 2D plane; shifting an image up by 5 pixels will result in the 2D convolution of that image shifting up by 5 pixels. In contrast, the number of atoms in a molecule is **invariant** under rotations and translations of the molecule in 3D space, and so the number of atoms is termed an SE(3) invariance. No matter how we rotate or translate the molecule, the number of atoms remains the same. In this work, we were able to achieve E(3) invariance (i.e. rotations, translations, and reflections) but not SE(3) invariance, since Euclidean distance matrices do not encode orientation. The mirror image of a chiral molecule is chemically distinct, however the E(3) invariant Euclidean distance matrices of our neural networks do not make this distinction.

### Shortcomings and future directions

While we desired an SE(3)-invariant network, we were only able to achieve E(3) invariance. Consequently, epRNA models predict an output which is defined only up to chirality, since chirality is not encoded in the distance representation. *Ad hoc* strategies to address this shortcoming include molecular dynamics simulation, which we performed in this work, or adding an additional chirality output from the network. Though we were able to create all-atom models using DMD, the RMSD was worse than for the epRNA prediction alone (Supplemental Table 1). The additional chirality output may be an extra sign which indicates if the positions produced by the multidimensional scaling should be mirrored in one dimension. Neither of these solutions are smoothly differentiable operations, and further work is needed to address the issue of chirality.

We omitted structures in PDB in which RNA is complexed with other macromolecules such as proteins, reducing the size of our training set. However, including such structures may introduce bias due to the presence of such interactions. Future researchers may focus on predicting interactions in protein-RNA complexes. The geometric methods in epRNA are widely applicable to macromolecule coordinate studies. We did not perform hyperparameter optimization; additionally, other architectures may also be feasible for RNA prediction, such as the commonly used transformer architecture (58, 59). Finally, we predicted only a single atom coordinate per backbone. Future work may extend epRNA to all-atom backbone prediction, particularly as more RNA structures are added to the Protein Data Bank.

### Potential applications

The epRNA approach may be useful in data in multimodal structural determination of RNAs, such as in NMR spectral analysis (60) or RNA sequencing after chemical cross-linking (61). It may also aid in rational design of RNA structures, by proposing approximate structures for given nucleotide sequences. Since epRNA inference is rapid and efficient, Monte Carlo design of RNAs can be accelerated. For example, if we wish to find an RNA sequence which folds into a desired 3D shape, we may begin with a single RNA base, propose moves (adding, removing, or changing a given residue) and accept or reject the move based on the RMSD of the epRNA predicted structure to the desired 3D shape. More generally, the geometric methods in the epRNA framework are widely applicable to molecular structure questions in biology and chemistry.

### Conclusion

We introduce a framework, epRNA, for direct prediction of RNA 3D structure from solely the nucleotide sequence. epRNA takes advantage of a smooth parametrization of Euclidean distance matrices, achieving rapid, robust, and accurate prediction of RNA 3D backbone structure across a range of nucleotide sizes. The epRNA framework addresses significant shortcomings in computational RNA structural biology, which has traditionally been limited by the speed of molecular dynamics simulations. By reasoning with the mathematical symmetries underlying the data, we achieve state-of-the-art predictions in RNA structure despite the limited structural data available for RNAs. We believe that the epRNA framework will be highly valuable to the structural biology and rational RNA engineering communities, and that such geometric approaches may be useful in other macromolecular structure problems.

## AUTHOR CONTRIBUTIONS

CMS implemented, trained, and validated the neural networks, and created the figures and wrote the initial draft of the manuscript. All authors conceptualized and discussed the ideas of this work.

## DATA AND CODE AVAILABILITY

We make our data and code publicly available on BitBucket (https://bitbucket.org/dokhlab/eprna-euclidean-parametrization-of-rna/src/master/).

## Abbreviations

epRNA: (Euclidean Parametrization of RNA)
PDB: (Protein Data Bank)
nts: (nucleotides)
EDM: (Euclidean distance matrix)
DMD: (discrete molecular dynamics)
NAST: (The Nucleic Acid Simulation Tool)
RMSD: (root mean square distance)

