## Supplemental Figures and Tables for "Predicting 3D RNA structure from solely the nucleotide sequence using Euclidean distance neural networks"

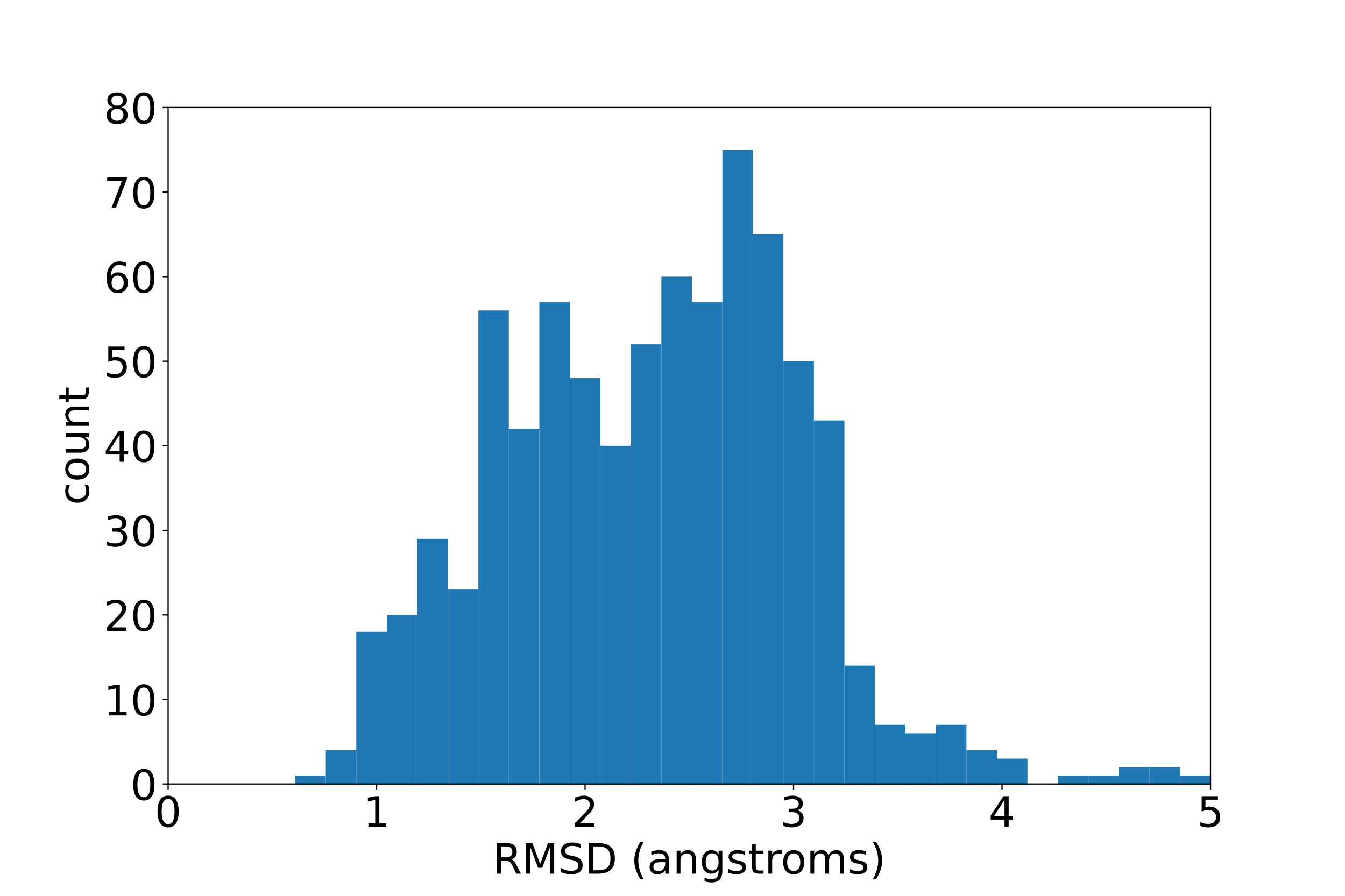

Supplemental Figure 1: Distribution of RMSDs (median: 2.4 angstroms, interquartile range: 1.05 angstroms) of RNA structures (n=788) taken from the Protein Data Bank.

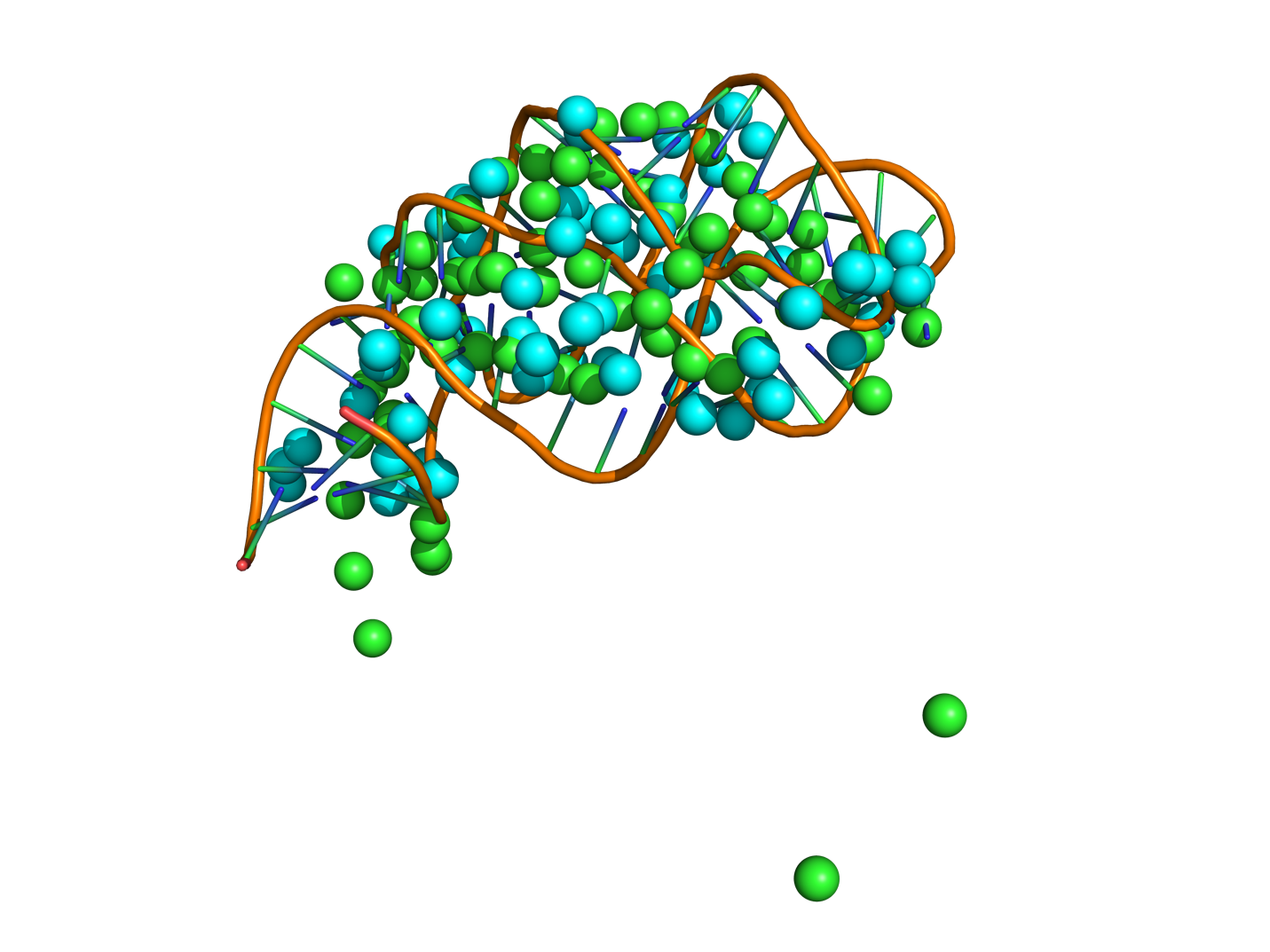

Supplemental Figure 2: 4FEO (orange) along with the original epRNA coordinates (green, 10 angstrom RMSD) and the NAST result (cyan, 18 angstrom RMSD).

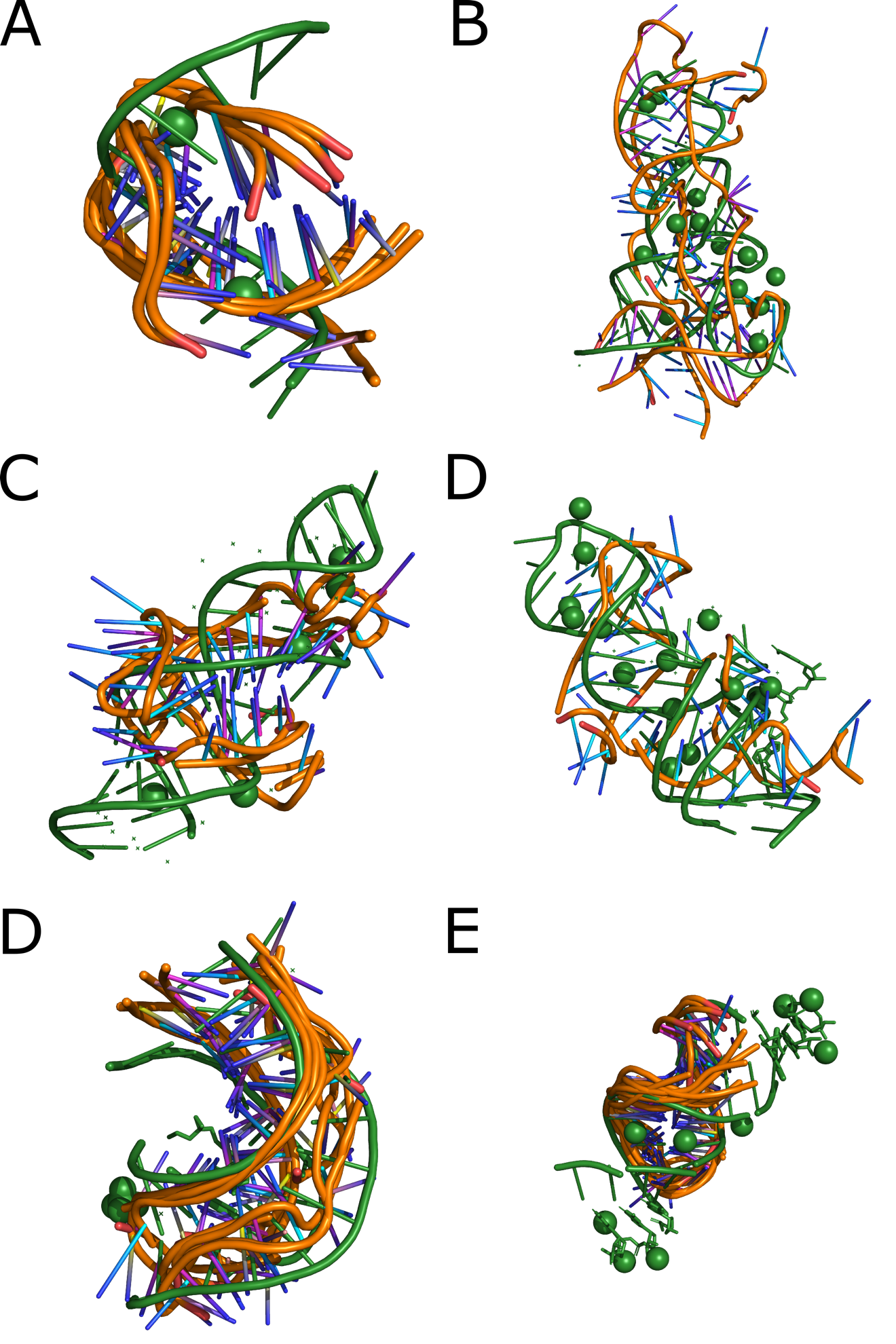

Supplemental Figure 3: DMD simulations (orange) compared to true crystal structures (green). **A-F**: 2Q1R, 3E5F, 4FNJ, 7D82, 7E9I, 7LNF.

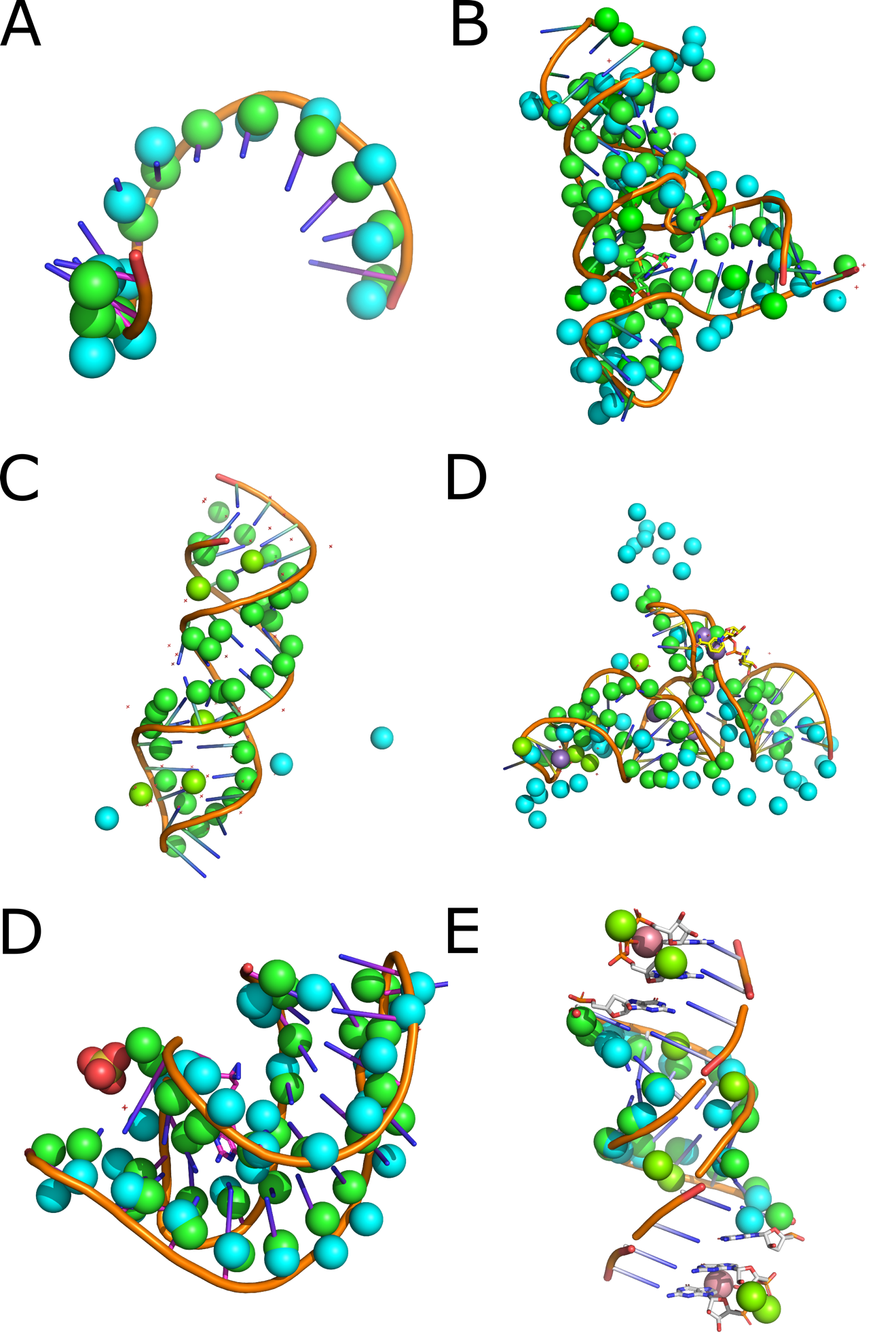

Supplemental Figure 4: NAST simulation results for the backbone (cyan) starting from the epRNA predicted configuration (green) overlaid with the true crystal structure (orange), with the same order of structures as in Supplemental Figure 3. Non-RNA atoms were not considered in simulations. NAST completely failed to converge in **C**, giving an RMSD >100 angstroms.

**Supplemental Tables**

Supplemental Table 1: Structural characteristics and results from molecular force field refinement of epRNA predicted structures. *p<0.05, **p<0.01, ***p<0.001

| **PDB ID** | **Length (number of nucleotides)** | **epRNA RMSD (angstroms)** | **Discrete molecular dynamics RMSD (angstroms)** | | **NAST RMSD (angstroms)** |
| --- | --- | --- | --- | --- | --- |
| 2Q1R | 12 | 1.5* | | 3.1 | 2.5 |
| 3E5F | 52 | 4.8*** | | 13.1*** | 8.0*** |
| 4FEO | 67 | 10.3*** | | 7.1*** | 18.6** |
| 4FNJ | 35 | 6.0*** | | 14.8* | >100 |
| 7D82 | 50 | 9.6*** | | 12.1*** | 16.7 |
| 7E9I | 33 | 2.1*** | | 3.8*** | 3.6*** |
| 7LNF | 18 | 1.2*** | | 3.8** | 2.8*** |
